# Unidirectional expression of enhancers with cell type-dependent direction of transcription

**DOI:** 10.1101/2023.06.27.546647

**Authors:** Saumya Agrawal, Emi Kanamaru, Yoriko Saito, Fumihiko Ishikawa, Michiel de Hoon

## Abstract

Enhancers are genomic regulatory elements that can affect expression of genes over megabases of genomic distances. As previous evidence suggested that transcription initiation at enhancers occurs in a bidirectional pattern, balanced expression of divergently transcribed capped RNAs has been used as a hallmark for enhancer detection from transcriptome data.

By analyzing deep Cap Analysis Gene Expression (CAGE) data from FANTOM5, FANTOM6, and other studies, we show that enhancers are usually unidirectionally transcribed in a given cell type. However, since the preferred direction of transcription of an enhancer may switch between cell types, enhancers may appear to be bidirectionally expressed when using CAGE data aggregated over many cell types. By analyzing expression directionality of enhancers across cell types, we classified enhancers into bidirectionally expressed, unidirectionally expressed with a consistent direction of transcription, and unidirectionally expressed with a switching direction, which was the largest category.

We conclude that requiring bidirectional expression during enhancer prediction from transcriptome data may lead to false negatives. Also, the preference of a given cell type for a specific direction of transcription at an enhancer suggests that enhancer RNAs are not transcriptional noise, but may be functional.

## Introduction

Enhancers are genomic regulatory elements that can affect expression of genes over megabases of genomic distances by chromatin looping [1, 2]. Enhancers produce capped transcripts known as enhancer RNAs (eRNAs) in a cell type-specific manner, with enhancer transcription associated with the regulatory activity of the enhancer [3, 4].

As eRNA transcription has been observed to occur in a bidirectional fashion [3], a bidirectional transcription pattern has been used as a signature feature for genome-wide enhancer identification [4, 5]. However, as enhancer RNAs are rapidly degraded by the exosome [6], their abundance is typically low, making the directionality of enhancer expression difficult to assess.

Here, we analyze enhancer expression directionality in recent deep sequenced CAGE (Cap Analysis Gene Expression) data sets. We find that enhancers are predominantly unidirectionally expressed in a given cell type. By analyzing enhancer directionality in 808 FANTOM5 CAGE libraries, we found preferential unidirectional transcription in most samples when analyzed separately, but bidirectional transcription when aggregating CAGE data across the libraries. In contrast to previous results, our analysis shows that most enhancers are unidirectionally expressed, with the preferred direction of transcription at each enhancer dependent on cell type.

## Results

We evaluate the expression directionality of each enhancer by calculating the Directionality Index [3]:

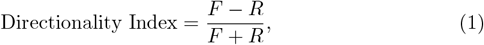

where *F* and *R* are the number of CAGE tags on the forward and reverse strand, respectively, at the enhancer. First, we calculated the Directionality Index for enhancers using pooled FANTOM5 Phase 1 CAGE data representing 808 cell types and tissues [7]. The distribution of the Directionality Index across enhancers is concave, showing that bidirectional enhancer expression is dominant (Figure 1A), in agreement with the previous analysis [4]. In contrast, using deep sequenced CAGE data from dermal fibroblast [8] (Figure 1B), from iPS [9] (Figure 1C), from a differentiation time course of the THP-1 leukemia cell line [10] (Figure 1D), and from human acute myeloid leukemia (AML) samples [11] (Figure 1E), we found a convex distribution of the Directionality Index, suggesting predominantly unidirectional enhancer expression.

**Figure 1.**
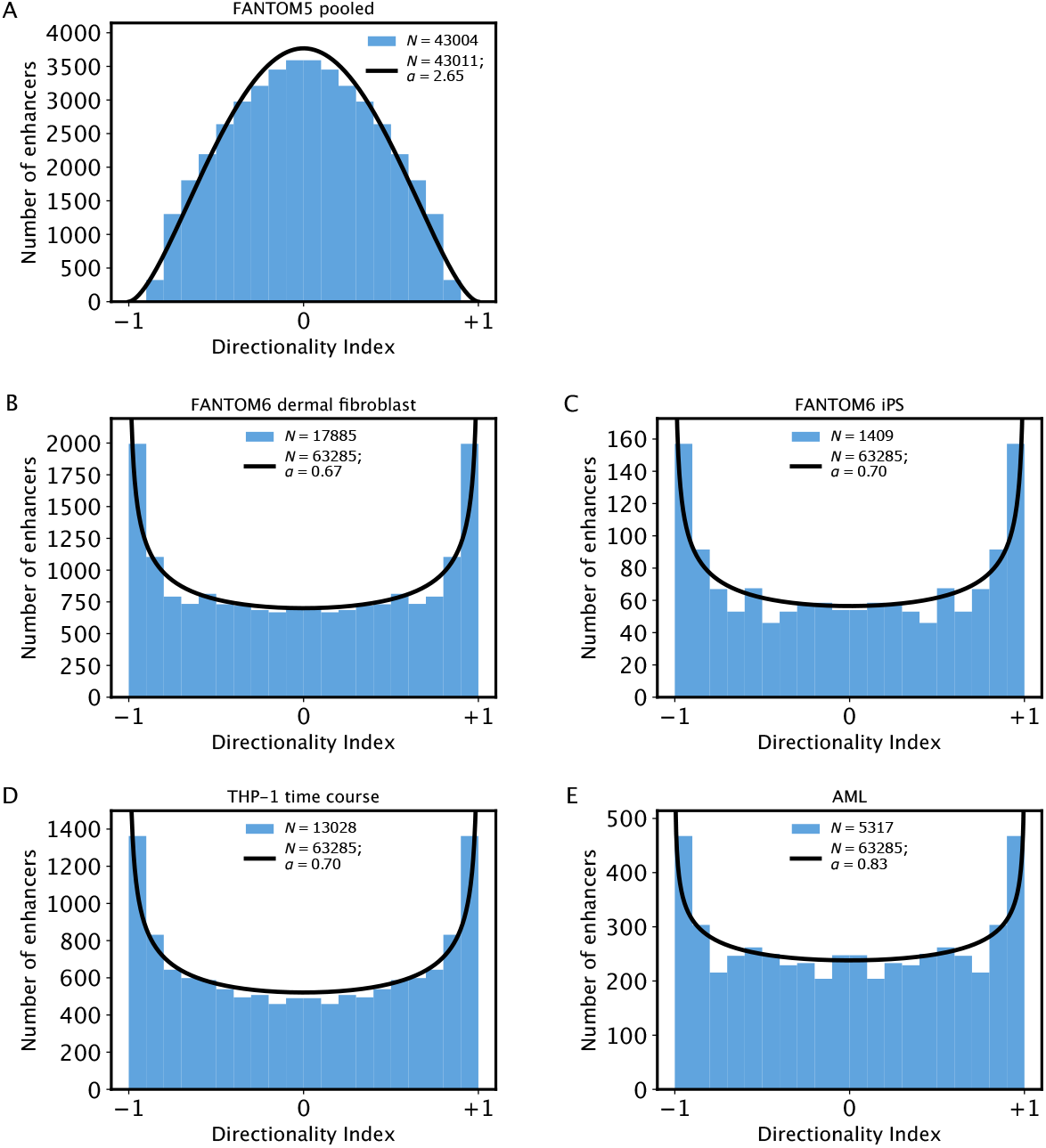
Distribution of the Directionality Index in (A) FANTOM5 pooled data, (B) FANTOM6 dermal fibroblast, (C) FANTOM6 iPS, (D) THP-1 time course, (E) AML. The estimated beta distribution is shown in black; the histogram calculated from enhancers with at least 10 CAGE tags is shown in blue. The number of enhancers *N* included in the calculation is shown in the legend of each panel; note that panel A uses the FANTOM5 Phase 1 enhancer set for human genome hg19, while panels B-E use the FANTOM5 Phase 1 + 2 enhancer set for human genome hg38.

A convex distribution (representing a preference for unidirectional enhancer expression) was obtained for CAGE data sets of a single cell type, while the distribution was concave (representing a preference for bidirectional expression of enhancers) for the FANTOM5 CAGE data, which were pooled across samples (421 from primary cell types, 252 from cell lines, and 135 from tissues) in the FANTOM5 Phase 1 collection. We then asked whether the distribution of the Directionality Index is concave or convex in each of the FANTOM5 samples separately. However, due to the limited sequencing depth in each of the samples, few enhancers have sufficient CAGE tags to estimate the Directionality Index accurately, precluding a direct calculation of a histogram.

Instead, we estimate the shape of the Directionality Index distribution by modeling it using a beta distribution. A beta distribution has two shape parameters, *α* and *β*. We set *β* = *α* to ensure that the distribution is symmetric, as the forward and reverse strand are defined arbitrarily for each chromosome. The symmetric beta distribution is convex if *α* = *β <* 1, concave if *α* = *β >* 1, and uniform over its domain if *α* = *β* = 1 (Figure 2).

**Figure 2.**
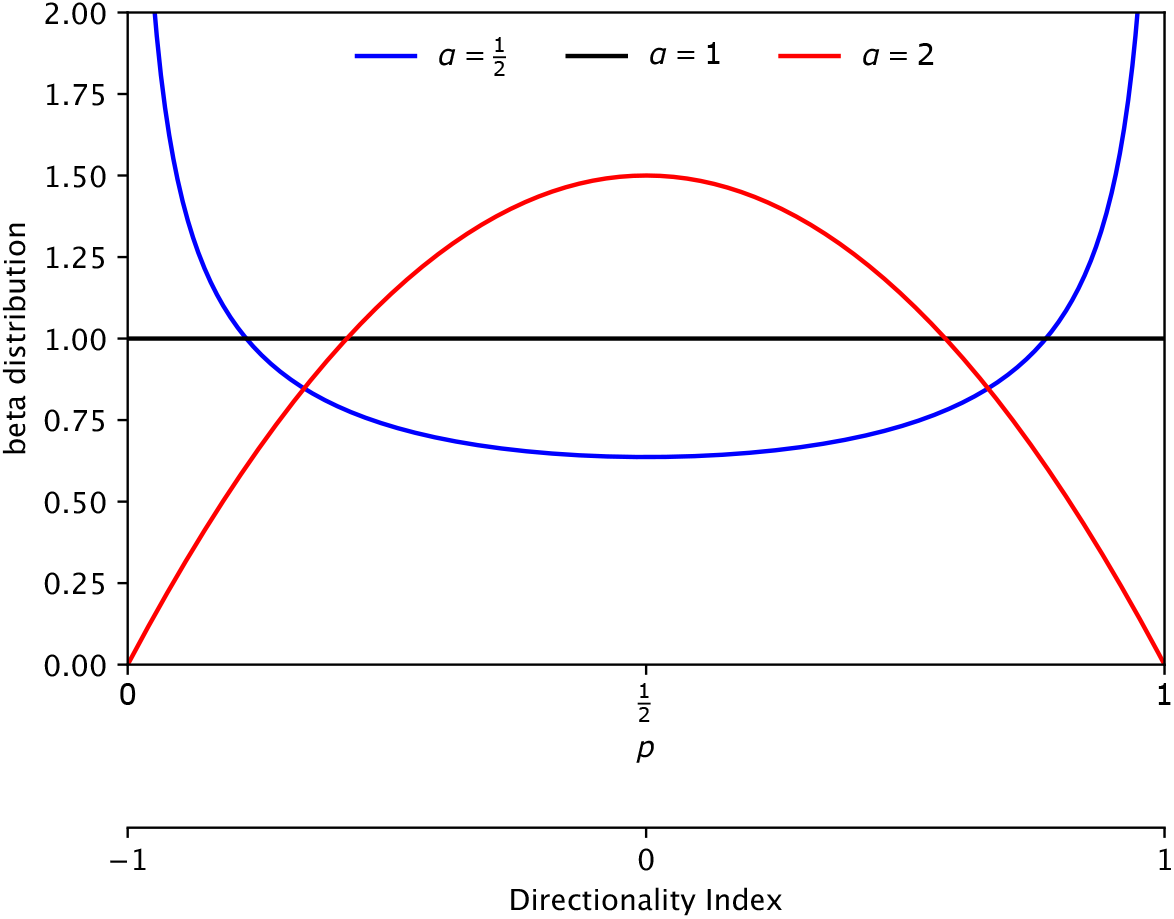
The beta distribution for *α* = *β <* 1 (convex), *α* = *β* = 1 (uniform), and *α* = *β >* 1 (concave).

For a given number of CAGE tags at a specific enhancer, we model the number of forward and reverse CAGE tags using the binomial distribution, where *p* is the probability for a CAGE tag to be on the forward strand. By compounding the beta and the binomial distribution, we find the beta-binomial distribution representing the number of forward and reverse CAGE tags at each enhancer (see Methods for details).

For the FANTOM5 Phase 1 data pooled over 808 samples, we found a shape parameter *α* = 2.65, which was significantly (*p <* 10^*−*100^, likelihood ratio test) greater than 1, consistent with bidirectional enhancer expression (Figure 1A). In contrast, we found a shape parameter *α* significantly smaller than 1 for the other CAGE data sets: dermal fibroblast (*α* = 0.67; *p <* 10^*−*100^), iPS (*α* = 0.70; *p* = 4 × 10^*−*42^), THP-1 (*α* = 0.70; *p <* 10^*−*100^), and AML (*α* = 0.83; *p* = 1.4×10^*−*31^) (likelihood ratio test; Figure 1B-E). These results are consistent with the pattern observed in the calculated histograms (Figure 1A-E).

Next, we applied the beta-binomial model to each of the 808 samples in the FANTOM5 Phase 1 CAGE data set separately (Figure 3). For 508 samples, the shape parameter was significantly (*p <* 0.05) smaller than 1, indicating significantly unidirectional enhancer expression. For 95 samples, the shape parameter was significantly (*p <* 0.05) greater than 1, indicating significantly bidirectional enhancer expression. For the remaining 205, the shape parameter was not significantly different from 1. We conclude that enhancer expression is predominantly unidirectional for most samples in the FANTOM5 Phase 1 data, while bidirectional enhancer expression is less common.

**Figure 3.**
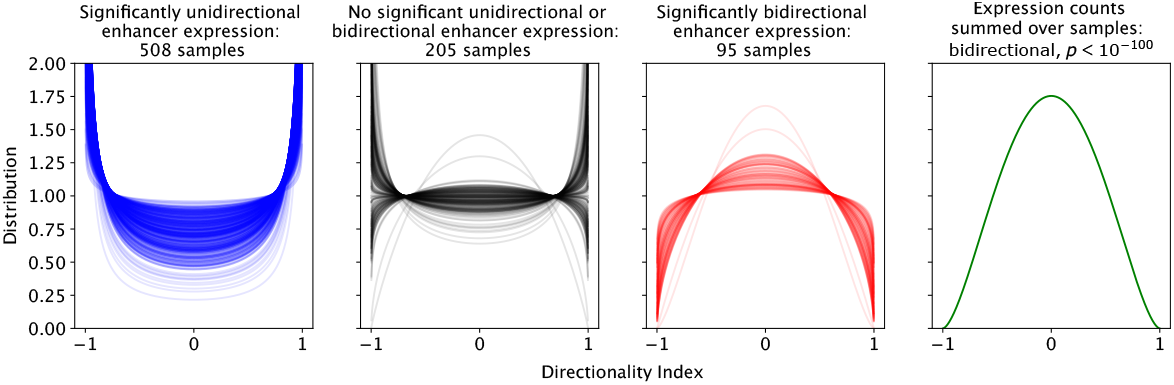
Distribution of the Directionality Index estimated for each sample in the FANTOM_5_ Phase 1 CAGE data separately.

Enhancer expression being predominantly unidirectional in individual samples but bidirectional in pooled data suggests that the preferred direction of expression of an enhancer can switch between samples. We modeled switching of the direction of enhancer expression by a modified binomial distribution

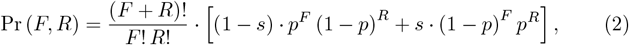

in which *s* is the switching probability. We apply this model to each enhancer by first setting *s* = 0 and performing the likelihood-ratio test to evaluate if *p* was significantly different from 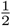 (corresponding to a Directionality Index of 0, i.e. perfectly balanced forward and reverse transcription). We classified the enhancer as bidirectionally expressed if *p* was not significantly different from 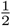. For *p* significantly different from 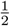, we next performed the likelihood-ratio test to evaluate if *s* was significantly different from 0. If it was, we classified the enhancer as switching; otherwise, we classified it as consistently unidirectionally expressed.

Applying this model to the FANTOM5 Phase 1 pooled CAGE data showed that 14,461 enhancers were unidirectionally expressed, with the same preferred direction of expression in all samples; 12,706 enhancers were bidirectionally expressed; and 15,844 enhancers were unidirectionally expressed, with the preferred direction of transcription switching between samples. As the sample collection in FANTOM5 does not comprise all human cell types, some of the enhancers classified as unidirectionally expressed may have a different preferred direction of expression in missing cell types, suggesting that the number of switching enhancers is an underestimate.

Using the FANTOM5 Phase 1 CAGE data, we find that switching enhancers form the largest category. We note that classification as switching enhancers is most stringent, as it requires reaching significance in two statistical tests (on *p* and *s*), while unidirectional enhancers require significance in only a single test (on *p*), and enhancers are classified as bidirectional if no statistical significance is found. Limitations in sequencing depth may therefore cause us to fail to identify unidirectional and switching enhancers, and overestimate the number of bidirectionally expressed enhancers.

Next, we searched for the human initiator motif [12] on both strands of each enhancer. The difference between the maximum score on the two strands was statistically significant for unidirectionally transcribed enhancers (*p* = 3.7 × 10^*−*10^, Mann-Whitney U test) but not for bidirectionally transcribed enhancers (*p* = 0.68, Mann-Whitney U test); the difference was marginally significant for switching enhancers (*p* = 0.0036, Mann-Whitney U test). Consistent with these results, the difference between the scores on the two strands was significantly larger for unidirectional enhancers compared to bidirectional enhancers (*p* =3.4 *×* 10^*−*5^).

## Discussion

Unidirectional [13, 14, 15, 16] and asymmetric [17, 18] expression of enhancers has occasionally been observed. In Drosophila, unidirectionally transcribed enhancers had an initiator (INR) motif for the transcribed strand only, while bidirectionally transcribed enhancers had an INR motif on both strands [19], suggesting that unidirectional or bidirectional transcription of enhancers is encoded in the genome sequence. In human, we also find a significantly higher INR motif score on the transcribed strand of unidirectional enhancers, while for bidirectional enhancers the two strands do not have a significantly different INR motif score.

While enhancer expression is indicative of enhancer activity [4], it remains unclear if enhancer transcription is due to transcriptional noise [20] or plays a biological role. For example, enhancer transcription may affect the local chromatin structure by displacing nucleosomes, may prevent gene silencing, or may stimulate the deposition of active chromatin marks [21]. Additionally, transcription factor binding to enhancer RNAs may boost the local transcription factor concentration and boost their regulatory effect [22]. The direction of enhancer transcription would modulate each of these mechanisms, providing an additional layer of regulation by enhancers.

As previous evidence indicated bidirectional transcription of enhancers, balanced expression of enhancer RNAs has been used as a criterion for enhancer detection from transcriptome data [4]. Our analysis shows that while enhancers are capable of bidirectional transcription, in any given cell type an enhancer may have a preferred direction of transcription. In particular when analyzing deep sequenced CAGE libraries obtained from one specific cell type, we recommend dropping bidirectional expression as a requirement for enhancer detection, as it may inadvertently discard enhancers that happen to be unidirectionally expressed in the given cell type.

## Methods

We use the binomial distribution to calculate the probability to have *F* forward CAGE tags and *R* reverse CAGE tags, for a given number of *F* + *R* CAGE tags in total:

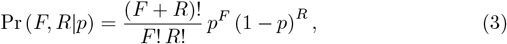

where *p* is the probability for a CAGE tag to be in the forward direction, and (1−*p*) is the probability for a CAGE tag to be in the reverse direction. With this parameterization, the expected value of the Directionality Index is 2*p* −1.

We model the distribution of *p* across enhancers using the beta distribution, with the parameters *α* and *β* chosen equal to each other such that the distribution is symmetric around 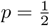:

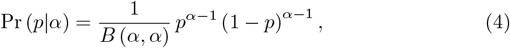

where *B* is the Beta function. The compound distribution is a beta-binomial distribution:

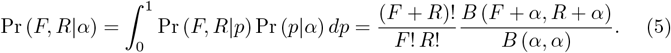

Dropping constant terms, the log-likelihood function for *n* enhancers is

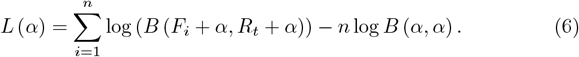

The parameter *α* is estimated using the maximum-likelihood method; its statistical significance is calculated using the likelihood-ratio test by comparing *L*(*α*) to *L*(*α* = 1).

To identify switching enhancers, we first set *s* = 0 and find *p* by maximizing the log-likelihood for the probability distribution given in Eq. (2):

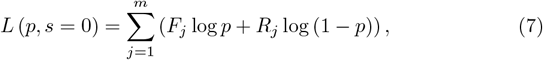

where *F*_*j*_ and *R*_*j*_ are the forward and reverse CAGE tag counts of the enhancer in sample *j*, and *m* is the number of samples. We compare *L* (*p, s* = 0) to *L*(*p* = 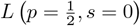 to evaluate if *p* is significantly different from 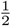. If it is, we next maximize the log-likelihood function

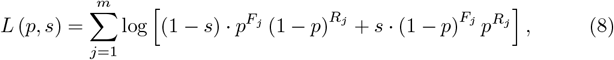

and use the likelihood-ratio test comparing *L* (*p, s*) to *L* (*p, s* = 0) to assess if the switching probability *s* is significantly different from 0.

## Acknowledgments

This work was supported by a Research Grant from MEXT to the RIKEN Center for Integrative Medical Sciences.

## References

[1] 1A. Oudelaar and D. Higgs, “The relationship between genome structure and function.” Nature Reviews Genetics 22: 154–168 (2021).

[2] H. Ray-Jones and M. Spivakov, “Transcriptional enhancers and their communication with gene promoters.” Cellular and Molecular Life Sciences 78: 6453–6485 (2021).

[3] T. Kim, M. Hemberg, J. Gray, A. Costa, D. Bear, J. Wu, D. Harmin, M. Laptewicz, K. Barbara-Haley, S. Kuersten, et al., “Widespread transcription at neuronal activity-regulated enhancers.” Nature 465: 182–187 (2010).

[4] R. Andersson, C. Gebhard, I. Miguel-Escalada, I. Hoof, J. Bornholdt, M. Boyd, Y. Chen, X. Zhao, C. Schmidl, T. Suzuki, et al., “An atlas of active enhancers across human cell types and tissues.” Nature 507: 455–461 (2014).

[5] N. Tippens, J. Liang, A. Leung, S. Wierbowski, A. Ozer, J. Booth, J. Lis, and H. Yu, “Transcription imparts architecture, function and logic to enhancer units.” Nature Genetics 52: 1067–1075 (2020).

[6] T. Hou and W. Kraus, “Spirits in the material world: Enhancer RNAs in transcriptional regulation.” Trends in Biochemical Sciences 46: 138–153 (2021).

[7] A. Forrest, H. Kawaji, M. Rehli, J. Baillie, M. de Hoon, V. Haberle, T. Lassmann, I. Kulakovskiy, M. Lizio, M. Itoh, et al., “A promoter-level mammalian expression atlas.” Nature 507: 462–470 (2014).

[8] J. Ramilowski, C. Yip, S. Agrawal, J. Chang, Y. Ciani, I. Kulakovskiy, M. Mendez, J. Ooi, J. Ouyang, N. Parkinson, et al., “Functional annotation of human long noncoding RNAs via molecular phenotyping.” Genome Research 30: 1060–1072 (2020).

[9] C. Yip, C. Hon, K. Yasuzawa, D. Sivaraman, J. Ramilowski, Y. Shibayama, S. Agrawal, A. Prabhu, C. Parr, J. Severin, et al., “Antisenseoligonucleotide-mediated perturbation of long non-coding RNA reveals functional features in stem cells and across cell types.” Cell Reports 41: 111893 (2022).

[10] I. Găzová, L. Lefevre, S. Bush, S. Clohisey, E. Arner, M. de Hoon, J. Severin, L. van Duin, R. Andersson, A. Lengeling, et al., “The transcriptional network that controls growth arrest and macrophage differentiation in the human myeloid leukemia cell line THP-1.” Frontiers in Cell and Developmental Biology 8: 498 (2020).

[11] M. Hashimoto, Y. Saito, R. Nakagawa, I. Ogahara, S. Takagi, S. Takata, H. Amitani, M. Endo, H. Yuki, J. Ramilowski, et al., “Combined inhibition of XIAP and BCL2 drives maximal therapeutic efficacy in genetically diverse aggressive acute myeloid leukemia.” Nature Cancer 2: 340–356 (2021).

[12] L. V. Ngoc, C. Cassidy, C. Huang, S. Duttke, and J. Kadonaga, “The human initiator is a distinct and abundant element that is precisely positioned in focused core promoters.” Genes & Development 31: 6–11 (2017).

[13] Y. Ho, F. Elefant, S. Liebhaber, and N. Cooke, “Locus control region transcription plays an active role in long-range gene activation.” Molecular Cell 23: 365–375 (2006).

[14] A. Kim, H. Zhao, I. Ifrim, and A. Dean, “Beta-globin intergenic transcription and histone acetylation dependent on an enhancer.” Molecular and Cellular Biology 27: 2980–2986 (2007).

[15] F. D. Santa, I. Barozzi, F. Mietton, S. Ghisletti, S. Polletti, B. Tusi, H. Muller, J. Ragoussis, C. Wei, and G. Natoli, “A large fraction of extragenic RNA pol II transcription sites overlap enhancers.” PLoS Biology 8: e1000384 (2010).

[16] F. Koch, R. Fenouil, M. Gut, P. Cauchy, T. Albert, J. Zacarias-Cabeza, S. Spicuglia, A. de la Chapelle, M. Heidemann, C. Hintermair, et al., “Transcription initiation platforms and GTF recruitment at tissue-specific enhancers and promoters.” Nature Structural & Molecular Biology 18: 956–963 (2011).

[17] M. Kowalczyk, J. Hughes, D. Garrick, M. Lynch, J. Sharpe, J. Sloane-Stanley, S. McGowan, M. D. Gobbi, M. Hosseini, D. Vernimmen, et al., “Intragenic enhancers act as alternative promoters.” Molecular Cell 45: 447–458 (2012).

[18] R. Chen, T. Down, P. Stempor, Q. Chen, T. Egelhofer, L. Hillier, T. Jeffers, and J. Ahringer, “The landscape of RNA polymerase II transcription initiation in C. elegans reveals promoter and enhancer architectures.” Genome Research 23: 1339–1347 (2013).

[19] O. Mikhaylichenko, V. Bondarenko, D. Harnett, I. Schor, M. Males, R. Viales, and E. Furlong, “The degree of enhancer or promoter activity is reflected by the levels and directionality of eRNA transcription.” Genes & Development 32: 42–57 (2018).

[20] K. Struhl, “Transcriptional noise and the fidelity of initiation by RNA polymerase II.” Nature Structural & Molecular Biology 14: 103–105 (2007).

[21] A. Field and K. Adelman, “Evaluating enhancer function and transcription.” Annual Review of Biochemistry 89: 213–234 (2020).

[22] A. Sigova, B. Abraham, X. Ji, B. Molinie, N. Hannett, Y. Guo, M. Jangi, C. Giallourakis, P. Sharp, and R. Young, “Transcription factor trapping by RNA in gene regulatory elements.” Science 350: 978–981 (2015).

